# Buffet-Style Expression Factor-Adjusted Discovery Increases the Yield of Robust Expression Quantitative Trait Loci

**DOI:** 10.1101/028878

**Authors:** Peter Castaldi, Ma’en Obeidat, Eitan Halper-Stromberg, Andrew Lamb, Margaret Parker, Robert Chase, Vincent Carey, Ruth Tal-Singer, Edwin Silverman, Don Sin, Peter D Paré, Craig Hersh

**Affiliations:** Channing Division of Network Medicine, Brigham and Women’s Hospital, Boston; Division of General Internal Medicine and Primary Care, Brigham and Women’s Hospital, Boston; University of British Columbia Center for Heart Lung Innovation, St Paul’s Hospital, Vancouver, BC, Canada; Computational Biosciences Program, University of Colorado, Denver; GSK R&D, King of Prussia, Pennsylvania; Pulmonary and Critical Care Division, Brigham and Women’s Hospital and Harvard Medical School, Boston; Respiratory Division, Department of Medicine, University of British Columbia, Vancouver, BC, Canada

**Keywords:** gene expression, expression quantitative trait loci, genomics

## Abstract

Expression quantitative trait locus (eQTL) analysis relates genetic variation to gene expression, and it has been shown that power to detect eQTLs is substantially increased by adjustment for measures of expression variability derived from singular value decomposition-based procedures (referred to as expression factors, or EFs). A potential downside to this approach is that power will be reduced for eQTL that are correlated with one or more EFs, but these approaches are commonly used in human eQTL studies on the assumption that this risk is low for *cis* (i.e. local) eQTL associations. Using two independent blood eQTL datasets, we show that this assumption is incorrect and that, in fact, 10-25% of eQTL that are significant without adjustment for EFs are no longer detected after EF adjustment. In addition, the majority of these “lost” eQTLs replicate in independent data, indicating that they are not spurious associations. Thus, in the ideal case, EFs would be re-estimated for each eQTL association test, as has been suggested by others; however, this is computationally infeasible for large datasets with densely imputed genotype data. We propose an alternative, “buffet-style” approach in which a series of EF and non-EF eQTL analyses are performed and significant eQTL discoveries are collected across these analyses. We demonstrate that standard methods to control the false discovery rate perform similarly between the single EF and buffet-style approaches, and we provide biological support for eQTL discovered by this approach in terms of immune cell-type specific enhancer enrichment in Roadmap Epigenomics and ENCODE cell lines.

**Significance Statement**: Genetic differences between individuals cause disease through their effects on the function of cells and tissues. One of the important biological changes affected by genetic differences is the expression of genes, which can be identified with expression quantitative trait locus (eQTL) analysis. Here we explore the basic methods for performing eQTL analysis, and we identify some underappreciated negative impacts of commonly applied methods, and propose a practical solution to improve the ability to identify genetic differences that affect gene expression levels, thereby improving the ability to understand the biological causes of many common diseases.

## Introduction

Expression quantitative trait locus (eQTL) analysis relates genetic variation to gene expression for the purpose of characterizing genetic loci that regulate the expression of local (*cis*) or distant *(trans)* genes(1). It is standard practice to adjust for broad components of expression variability (referred to as “expression factors” or EFs), because this has been shown to dramatically increase the yield of discovered eQTL(2). This approach is effective because gene expression can be influenced by multiple sources, including biological processes, environmental exposures, and technical artifacts. Alter et al. first demonstrated the utility of applying principal components analysis (PCA) to gene expression data, but they also acknowledged that it is challenging to correctly identify and separate extraneous noise from the desired signal. Supervised methods were subsequently developed to identify factors that were largely independent from the primary signal of interest, including surrogate variable analysis (SVA)(3) and the probabilistic estimation of expression residuals (PEER)(4) method. While these widely used methods can eliminate technical artifacts, they may also remove true biologic signals which may or may not be relevant to the primary analysis.

Unlike standard differential expression analyses that test thousands of hypotheses, genomewide eQTL analyses test millions. Ideally, EFs would be re-estimated for each tested eQTL hypothesis (5, 6). A major limitation of these approaches is their computational expense, which may be prohibitive for genomewide imputed SNP data and increasingly detailed measurements of expression. Thus, it has been suggested that for local eQTL studies it is appropriate to estimate factors once and then use this set of factors for each association test(4, 5). The potential risk is that if EFs are correlated with an eQTL association of interest, these adjustments can decrease power. This limitation has been demonstrated for eQTL hotspots, i.e. genetic loci that regulate many genes in *trans*(7), and adjustment for EFs has been shown to decrease observed heritability of some gene expression levels (8).

While adjustment for EFs generally increases the overall yield of eQTL analyses, the fine-grained impact of these adjustments on eQTL discovery has not been well-characterized. We hypothesized that EF adjustment may decrease power to discover certain subsets of true local eQTL associations. To test this hypothesis and understand the impact of EF adjustment on eQTL analysis in more detail, we performed eQTL analyses including between 0-80 PEER factors from whole blood gene expression. To distinguish true from false eQTL discoveries, we performed replication analyses in a larger sample of separate subjects from the same study. We examine the merits of a more comprehensive yet computationally manageable approach for eQTL discovery (“buffet-style” eQTL discovery) that performs multiple PEER and non-PEER adjusted analyses to maximize the yield of eQTLs, rather than performing a single PEER-adjusted eQTL analysis.

## Results

Subjects with chronic obstructive pulmonary disease (COPD) were enrolled in the multicenter ECLIPSE Study(9). The demographic characteristics of the test (ECLIPSE 1) and replication (ECLIPSE 2) samples are similar (Supplemental Table 1). ECLIPSE 1 consisted entirely of former smokers, whereas 24% of the ECLIPSE 2 set were current smokers.

**Table 1.** Yield and Retention of Chromosome 15 eQTL Genes in PEER and non-PEER Adjusted Analyses in Two Data Sets

|  | ECLIPSE 1 |  |  | ECLIPSE 2 |  |  |
| --- | --- | --- | --- | --- | --- | --- |
|  | # PEER Factors | # eQTL Genes | % Baseline | # PEER Factors | # eQTL Genes | % Baseline |
| Baseline | NA | 165 | NA | NA | 273 | NA |
| Max Yield PEER | 24 | 271 | 74 | 51 | 413 | 91 |
| PEER + Cell Count | 20 | 269 | 75 | 52 | 408 | 92 |
| Baseline eQTL analysis adjusts only for first 3 principal components of genetic ancestry.<br>Max Yield PEER analysis is the PEER-adjusted analysis with the highest yield of eQTL genes.<br># PEER factors is the number of adjusted PEER factors in the specified analysis.<br>PEER + Cell Count – analyses directly adjusted for white blood cell count percentages (neutrophils, lymphocytes, monocytes, and eosinophils).<br>% Baseline is the percentage of significant baseline eQTL genes that were also significant in the specified analysis.<br>For all analyses, significance was defined as q-value $\leq 0.05$. |  |  |  |  |  |  |

To assess the biological associations of PEER factors, we tested ECLIPSE 2 PEER factors for association with clinical covariates, environmental exposures, white blood cell differential counts and protein biomarkers. We identified multiple strong associations indicating that many PEER factors were related to biological variables (Figure 1).

**Figure 1.**
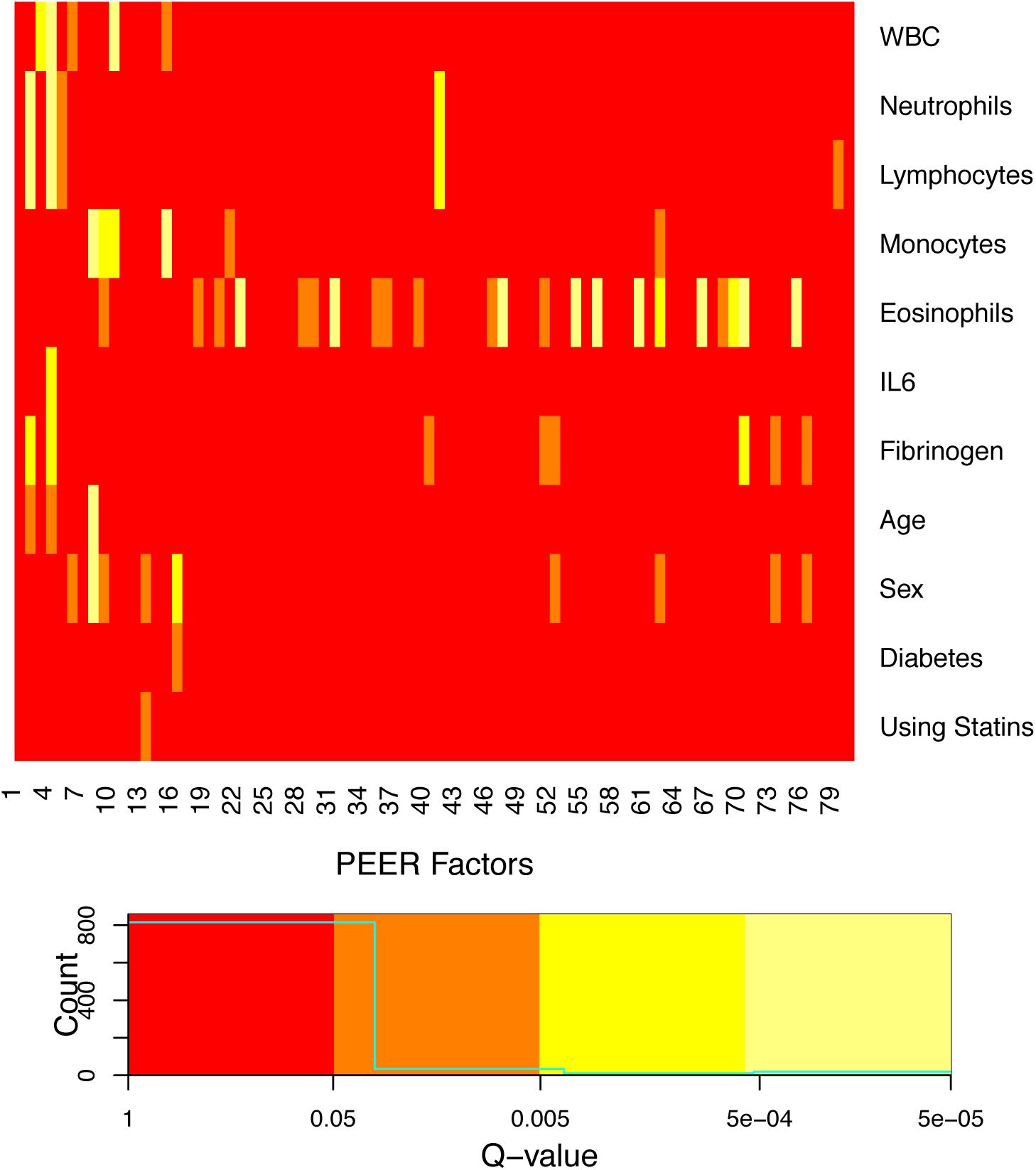
Heatmap of Associations Between PEER Factors, Cell Proportions, and Clinical Measures in ECLIPSE 2. PEER factors capture significant biological and clinicallyDinduced patterns in the expression data, and these biologic and clinical patterns are often represented in multiple PEER factors.

### Chromosome 15 Analysis

Due to the computational requirements of genome-wide eQTL analysis, initial analyses were conducted in chromosomes 15 and 22, with consistent results. The chromosome 22 results are shown in the Supplemental Materials.

Table 1 summarizes the results of the local eQTL (i.e. within 1MB of the gene boundaries) analyses for chromosome 15. Adjustment for PEER factors increased the yield of significant eQTL genes in both ECLIPSE 1 and 2 (Table 1 and Supplemental Table 2). Figure 2 shows the relationship between the number of PEER factors used for adjustment and the yield of local eQTL genes. In ECLIPSE 1 (panel A), a saturation point is reached at 24 PEER factors with diminishing returns with further adjustment. In ECLIPSE 2 (panel B), yield peaked at 51 PEER factors but did not appreciably decrease with further PEER adjustment. Comparison of the baseline eQTL analysis to the PEER-adjusted analysis yielding the largest number of eQTL genes (referred to subsequently as the *max-yield analysis*), showed a loss of approximately 10-25% of eQTL genes that were significant in the baseline analysis (Table 1). A separate analysis adjusting for both cell count differential in addition to PEER factors did not increase the yield of significant eQTL.

**Table 2.** Yield and Retention of eQTL Genes in PEER and non-PEER Adjusted Genome-wide eQTL Analyses in Two Data Sets

|  | ECLIPSE 1 |  |  | ECLIPSE 2 |  |  |
| --- | --- | --- | --- | --- | --- | --- |
|  | # PEER Factors | # eQTL Genes | % Baseline | # PEER Factors | # eQTL Genes | % Baseline |
| Baseline | NA | 4358 | NA | NA | 7295 | NA |
| Max Yield PEER | 15 | 7978 | 82 | 42 | 11632 | 90 |
| Buffet-Style | 15 | 11426 | 100* | 45 | 14438 | 100* |
| Baseline eQTL analysis adjusts only for first 3 principal components of genetic ancestry.<br>The maximum number of PEER factors used in eQTL analysis was 15 and 45 for ECLIPSE 1 and ECLIPSE 2, respectively.<br>Max Yield PEER analysis is the PEER-adjusted analysis with the highest yield of eQTL genes.<br>Buffet-Style analysis is an analysis that retains all eQTLs associations with a q-value $\leq 0.05$ in any of the PEER or non-PEER adjusted analyses.<br># PEER factors is the number of adjusted PEER factors in the specified analysis.<br>% Baseline is the percentage of significant baseline eQTL genes that were also significant in the specified analysis.<br>* 100% by definition, since buffet-style results include significant baseline associations.<br>For all analyses, significance was defined as q-value $\leq 0.05$. |  |  |  |  |  |  |

**Figure 2.**
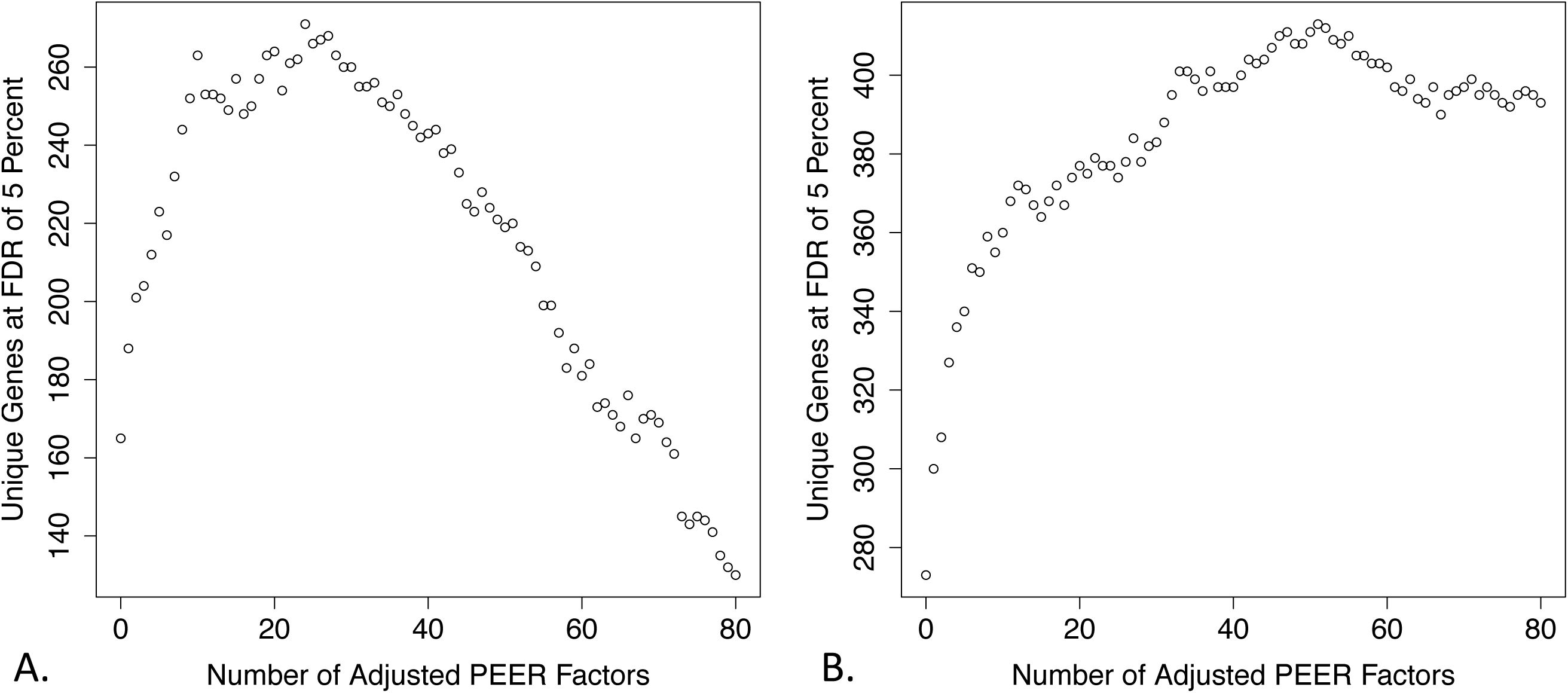
Discovery of eQTL Genes in Multiple PEER and non-PEER Adjusted Analyses in ECLIPSE 1 (Panel A) and ECLIPSE 2 (Panel B). For both ECLIPSE 1 and ECLIPSE 2, there is a large increase in the number of significant eQTL genes with adjustment for the first 10-20 PEER factors.

The replication rates (replication = the proportion of eQTL genes significant in the ECLIPSE 2 baseline or max yield analysis) for baseline and max-yield eQTL genes in ECLIPSE 1 were 85% and 81%, respectively demonstrating that these associations are not spurious due to technical artifacts. A comprehensive representation of the replication rates for all 81 analyses performed in ECLIPSE 1 and 2 shows that the highest number of replicated genes coincides with the number of factors producing the highest yield in ECLIPSE 1 (Figure 3, Panel A), and that replication rates are consistent across most analyses (Panel B). The lowest replication rate is observed for the non-PEER adjusted analyses in ECLIPSE 1 and 2, and the highest replication occurs for analyses adjusting for the largest number of PEER factors (Panel B). However, the modest increase in replication at this level of PEER adjustment comes at the cost of far fewer eQTL discoveries. When we examined a more stringent replication, i.e. the identical SNP-gene pair, the replication rates for the baseline and max yield ECLIPSE 1 analyses were 55% and 51%, respectively. Both were significantly higher than the 3% replication rate for a randomly selected set of non-significant SNP-gene pairs in ECLIPSE 1.

**Figure 3.**
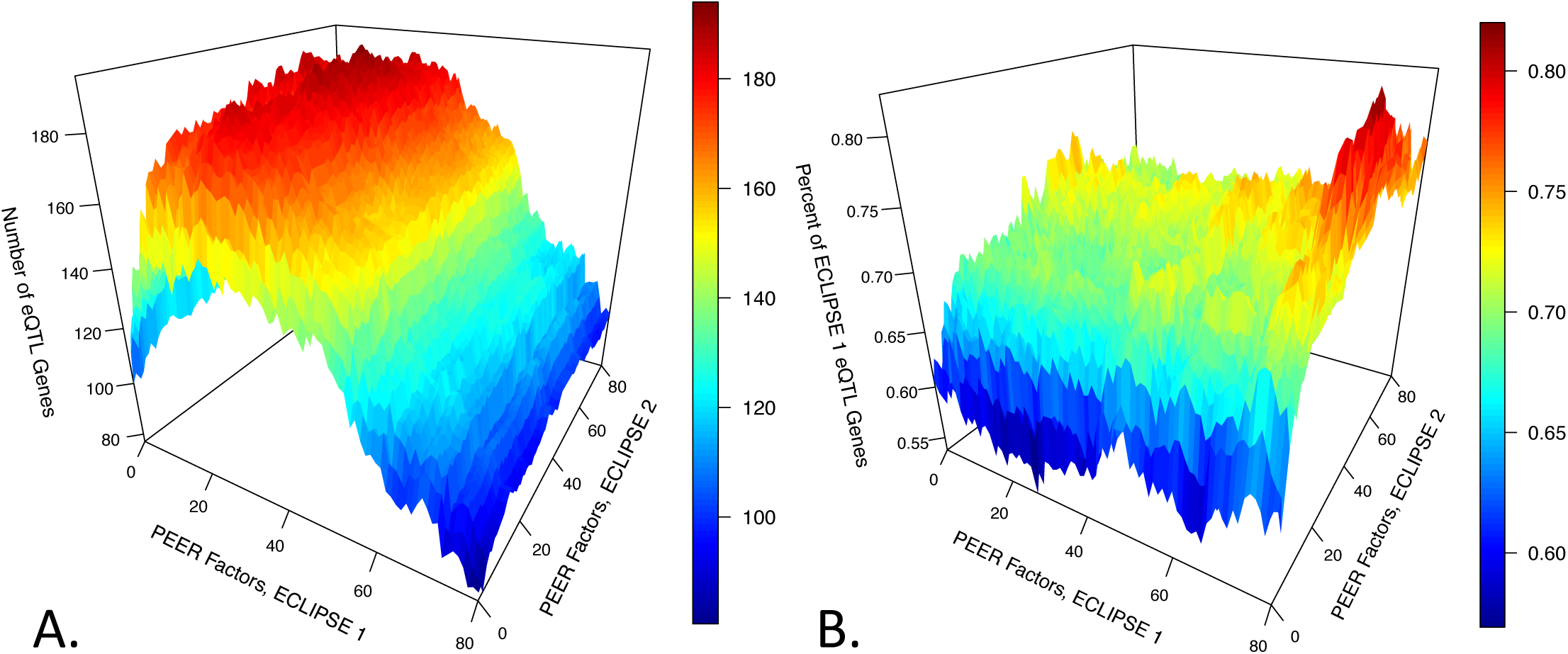
Independent Replication for eQTL Genes Identified in Multiple PEER and non-PEER Adjusted Analyses. For each combination of PEER adjusted analyses in ECLIPSE 1 and ECLIPSE 2, the number of consistent (i.e. replicated at q-value <0.05) eQTL genes (Panel A) and the replication rates (Panel B) are shown.

In the sequence of analyses starting with the baseline (non-PEER adjusted) analysis, adjustment for the first few PEER factors dramatically increases the yield of eQTL genes (Figure 4). While additional PEER adjustment yields a smaller number of novel eQTL genes (i.e., eQTL genes that had not achieved significance earlier in the sequence), such genes continue to be discovered even beyond the number of maximum yield PEER factors. While the common practice of focusing only on the max yield PEER analysis increases the number of discoveries, it also fails to identify a number of eQTL associations that would have been significant in an analysis adjusted for a different number of PEER factors or no PEER factors at all.

**Figure 4.**
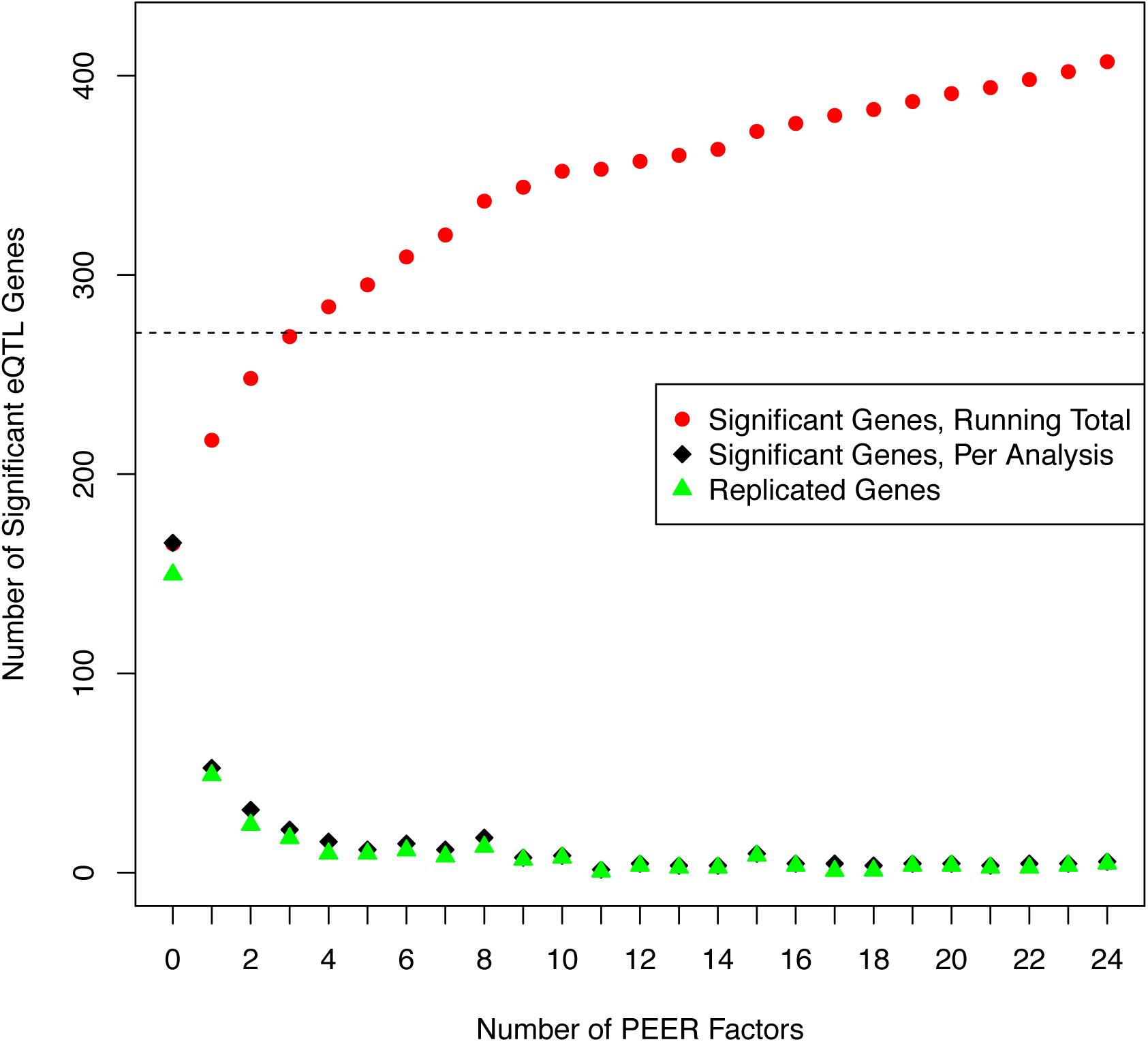
Cumulative Yield of eQTL Genes with PEER Adjustment. As the number of adjusted PEER factors is increased, the cumulative yield of eQTL from buffet-style collection of significant ECLIPSE 1 results is shown (red dots), as well as the number of significant novel genes in each analysis (black dots) and the number of novel genes that are also significant in ECLIPSE 2.

As an alternative to focusing only on the max yield PEER analysis, we propose a “buffet-style” approach in which all significant results at a 5% FDR threshold are collected from the series of eQTL analyses from the baseline up to adjustment for 15 PEER factors. This approach resulted in 372 significant eQTL genes on chromosome 15, compared to 271 significant genes obtained from the max yield ECLIPSE 1 analysis. Since these q-values were calculated only in the individual analyses and not the entire set of p-values, we compared the analysis-specific q-values to the q-values calculated from concatenating the ~76 million p-values from all sixteen analyses. The distributions of these two sets of q-values are nearly identical (Supplemental Figure 3). Furthermore, if the q-values calculated from all concatenated p-values are used to determine significance, the number of significant genes is 382 (overlap with 372 buffet-style significant genes = 369,99.2% concordance).

We also tested the performance of PEER and non-PEER adjusted eQTL discovery (0 to 15 PEER factors) using the permutation-based assessment of eQTL gene significance employed in the GTEx project(10). Supplemental Figure 4 shows that the impact of PEER adjustment on eQTL discovery was similar in the permutation-based approach, while the overall yield of genes was decreased across all analyses. Replication of eQTL genes identified from each of the 16 analyses ranged from 75-81%. The buffet-style approach identified 161 significant genes, compared to 119 significant genes identified from the max-yield analysis. All 161 significant buffet-style genes from the permutation-based approach were also present in the standard buffet-style results.

### Genome-wide Local eQTL Analysis

We repeated the above local eQTL analyses on a genome-wide level, using 15 PEER factors for ECLIPSE 1 and 45 PEER factors for ECLIPSE 2 (roughly ten observations per covariate). The effect of PEER adjustment was comparable to the chromosome 15 results (Table 2), with gene level replication rates of 81% and 73% and SNP-gene level replication rates of 55% and 51% for the baseline and max yield analyses, respectively. At the SNP-gene level, the same direction of effect for replicated eQTL was observed for 89-95% of replicated associations for the four possible combinations of baseline and max-yield analyses in ECLIPSE 1 and 2). The buffet-style discovery approach in ECLIPSE 1 led to a 43% increase in significant eQTL genes over the max yield analysis. The gene-based replication rates for all identified eQTL genes and “buffet-style only” genes (N=3348) not identified in the base or max-yield analyses were 77% and 72%, respectively. We compared the significant ECLIPSE 1 SNP-gene pairs from the baseline and max-yield analyses to the whole blood eQTL results from the GTEx project (v4), and we observed a 58% gene-level and 31% SNP-gene level replication rate.

In ECLIPSE1, the baseline only eQTL (n=803 unique genes) have larger effect sizes than “max-yield only” eQTL (n=4423 genes) (Supplemental Figure 5). Furthermore, the baseline-only eQTL showed lower q-values in the max-yield analysis (Supplemental Figure 6). Baseline only eQTL tend to have stronger effects and show more consistency across PEER and non-PEER adjusted analyses, whereas many max-yield only eQTL lie well below the detection limit of non-PEER adjusted analyses. In Figure 5, local association plots are shown for representative baseline only and max yield only eQTL for the *AKT3* and *SNX6* genes. In both instances, the lead eQTL SNP-gene association was replicated in ECLIPSE 2.

**Figure 5.**
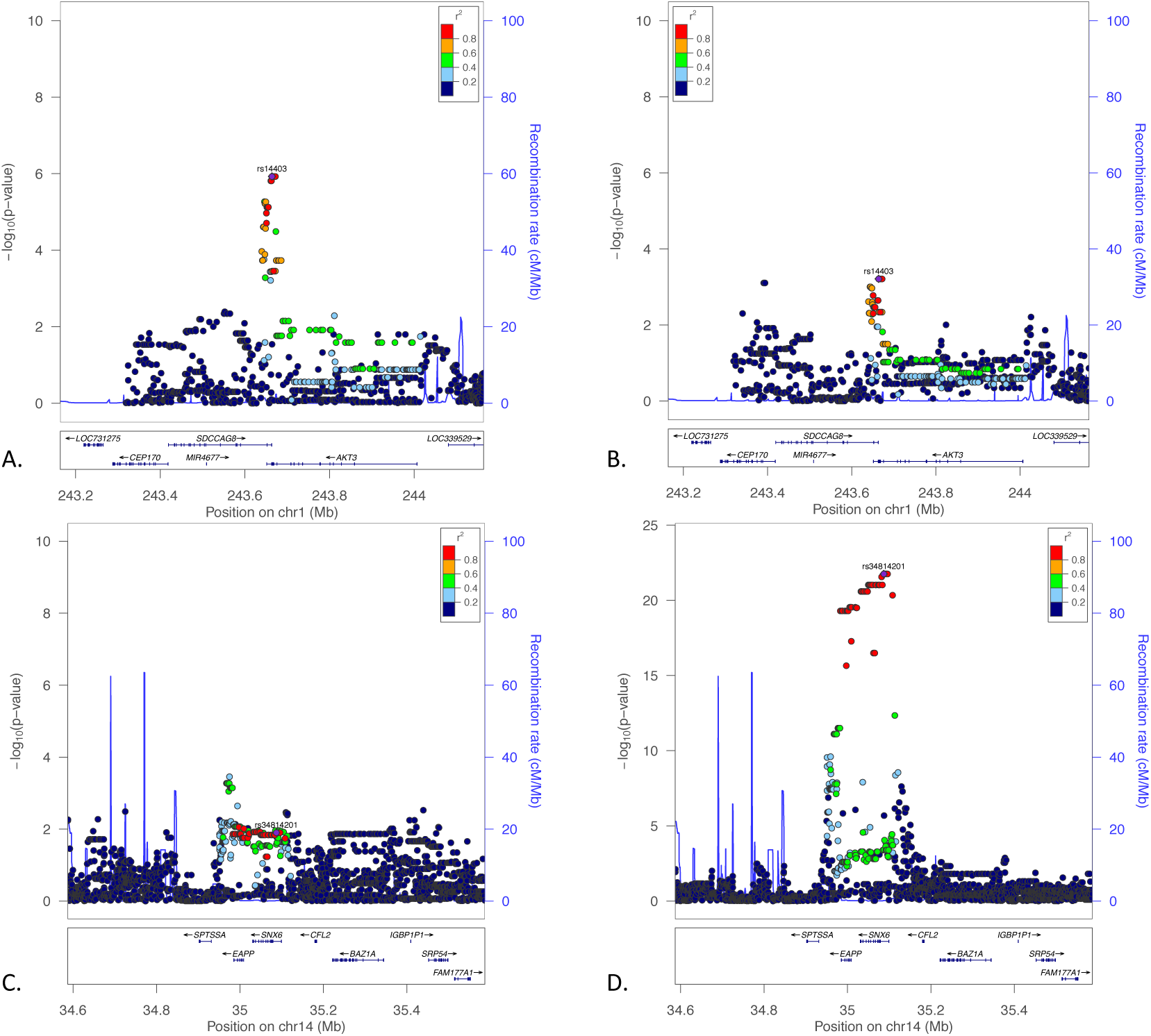
Local Association Plots for Baseline Only and Max-Yield Only eQTL Associations in *AKT3* and *SNX6.* In *AKT3,* the lead eQTL SNP exceeds the significance threshold in the baseline only (Panel A), but not the max-yield analysis (Panel B). Conversely, the lead eQTL SNP for *SNX6* is well below the significance threshold in the baseline analysis (Panel C), but very strongly associated with the PEER-adjusted expression of *SNX6* in the max-yield analysis (Panel D).

**Figure 6.**
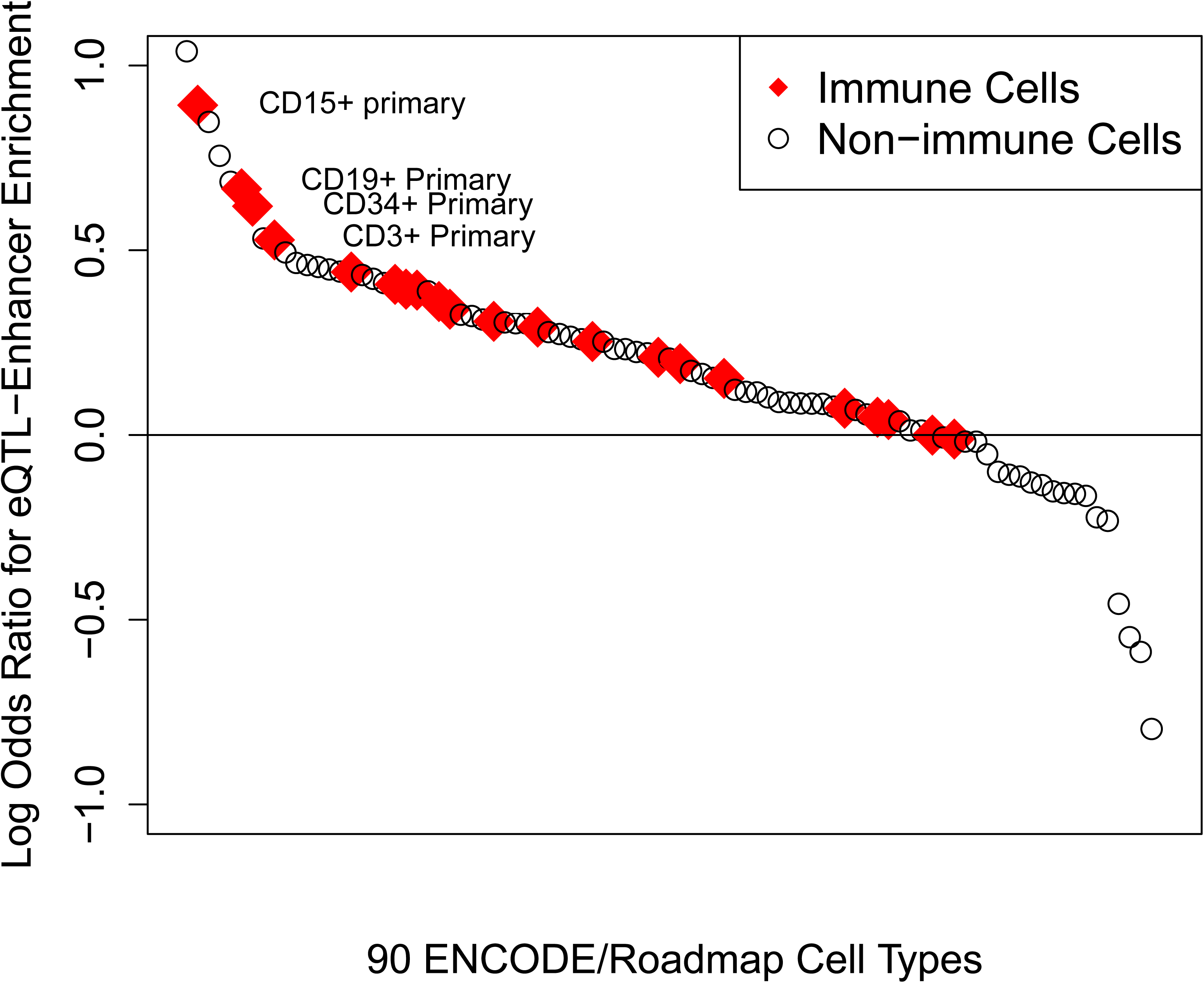
Cell Type-Specific Enhancer Enrichment for Top 100 Whole Blood eQTLs from Buffet-Style Analysis. Normalized measures of enrichment indicate that top blood eQTLs are enriched in enhancer regions from multiple cell types, with a preponderance of immune-related cell types showing the strongest enrichment (p=0.03, Mann-Whitney test).

We selected the lead SNPs from the 100 most strongly associated ECLIPSE 1 eQTL genes, and we used the probabilistic identification of causal SNPs (PICS) approach(11) to identify all SNPs estimated to have a >5% likelihood to be the causal variant for each gene. In the Roadmap Epigenomics data(12), we observed significant enrichment of these SNPs in enhancer regions of immune-related cell types (n=21) versus all other cell types (n=69, p=0.03), consistent with the cells found in whole blood (Figure 6). For eQTL discovered by the buffet-style approach but not in the max-yield analysis we observed a similar pattern of strong enrichment for immune-related cell types (p<0.001, Supplemental Figure 7).

## Discussion

The complex relationship between EFs (i.e. expression PCs or PEER factors), technical artifacts in expression data, and biological signal is well established(5,10). The common practice of estimating factors once rather than iteratively with each SNP-gene contrast and then using these as covariates for genomewide eQTL discovery rests on the assumption that these factors are independent of true SNP-gene associations. While this is known to be false for distant eQTL hotspots, there is conflicting evidence on the validity of this assumption for local eQTLs. Our data suggest that these adjustments reduce power for some eQTLs. We present the following main findings: 1) Adjustment for EFs simultaneously increases the overall eQTL yield while also resulting in the loss of some significant eQTL identified without EF adjustment. 2) Both EF-identified and non-EF identified eQTL genes show comparable rates of independent replication. 3) The complex effect of EF adjustment on eQTL discovery precludes the identification of an optimal number of EFs that can be uniformly applied across all tests in a given eQTL study.

Alter et al. first described the application of singular variable decomposition (SVD) to expression arrays to summarize expression heterogeneity and identify biologically interpretable “eigengenes.” They acknowledge that the identification of the optimal rotation for such factors is uncertain, and they suggest selecting a rotation that maximizes the “biological interpretability” of the resulting factors(13). Storey and Leek subsequently developed surrogate variable analysis (SVA), a supervised approach that specifically considers the potential for correlation between EFs and the primary variable of interest in an expression study(3). This work was extended by others(7,14,15), with specific methods proposed for SNP-by-SNP estimation of EFs(5, 6). Fusi et al. developed the PANAMA method which simultaneously models the effect of a subset of genetic variants and EFs, though this approach was applied only to SNPs with large *trans* effects.(16). Most relevant to the issue of estimating EFs for local eQTL analysis, Stegle et al. implemented a Bayesian procedure that learns EFs separately for each genetic variant(5). While they demonstrate that this procedure is superior for the identification of local and distant eQTL in yeast data, they also suggest that for human cis eQTL data it may be reasonable to eschew the computational burden of iterative estimation of EFs. This PEER factor-based approach has been used in a number of eQTL studies, including GTEx(10). Goldinger et al. come to different conclusions based on a heritability approach that indicates that EFs often show significant heritability and can decrease the observed heritability of some gene expression levels(8).

To address this uncertainty, we performed multiple analyses adjusting for a wide range of EFs. The use of a large replication dataset provides a reasonable standard to distinguish true from false eQTL discoveries. Our data indicate that while EF adjustment is generally beneficial, it excludes an appreciable number of eQTL that would otherwise be identified in a non-adjusted analysis. Furthermore, these eQTL show reasonable replication rates, indicating that these are not due to spurious association. We also demonstrate that novel eQTL are discovered across a wide range of EFs. While the common practice of selecting the maximal yield EF adjusted analysis clearly increases the yield of eQTL over a non-adjusted analysis, it is apparent that this also excludes true eQTLs. The clear benefits of EF adjustment are associated with less-appreciated costs, and it is difficult to justify the choice of a single number of EFs to apply uniformly across a genome-wide eQTL analysis.

We propose a “buffet-style” approach in which significant eQTL are collected from multiple non-EF and EF adjusted analyses, which results in a notable increase in eQTL discoveries. This represents a compromise between SNP-by-SNP re-estimation of EFs and the current practice of estimating EFs once for a genome-wide analysis. We demonstrate that standard methods of FDR control behave similarly in the buffet-style approach compared to a single eQTL analysis, since the distribution of p-values across analyses is similar. However, the performance of various FDR-based methods under conditions of strong dependence (as is characteristic of densely imputed genotype data) remains an area of ongoing investigation(17). Permutation-based significance assessment was a more stringent approach yielding fewer significant associations, yet the pattern with respect to eQTL discovery and replication was the same.

This study provides a systematic view of the effect of EF adjustment on eQTL discovery, and the use of two independent data sets enables quality assessment of the different eQTL analytical options. This study used whole blood expression data, so it is not clear how these results will generalize to other tissues. However, it is likely these observed phenomena of EF adjustment are generic to eQTL analysis.

In summary, EF adjustment increases eQTL yield, but fails to detect a subset of valid eQTL. EF adjustment reasonably accounts for potential biological confounders such as cell differential, but the presence of biological signals in EFs also may reduce power for discovery of eQTL that are collinear with one or more EFs. EF adjustment is a powerful tool to augment eQTL discovery, but eQTL results are sensitive to the number of EFs. While the optimal approach to eQTL discovery is likely iterative re-estimation of EFs for each SNP-gene association, in the absence of a computationally feasible implementation a reasonable alternative is the buffet-style pooling of eQTL results across a range of EF and non-adjusted analyses.

## Methods

### Study Samples

ECLIPSE (Evaluation of COPD Longitudinally to Identify Predictive Surrogate Endpoints) is an observational study of current and former smokers. (9). For this analysis, we selected two non-overlapping subsets of 130 (referred to as ECLIPSE 1) and 490 subjects (referred to as ECLIPSE 2). White blood cell analysis was performed on blood drawn concurrently for PaxGene RNA samples.

### Gene Expression

Gene expression profiling was performed using the Affymetrix Human U133 Plus2 array in ECLIPSE 1 and the HumanGene ST 1.0 array in ECLIPSE 2. The HumanGene ST 1.0 array was analyzed at the gene level. Gene expression data were log-transformed, and background correction was performed using robust multi-array averaging (RMA) (18). The ECLIPSE 2 data were generated in multiple batches, and ComBat was used to remove batch effect(19). After quantile normalization, expression values were rank transformed (20). For each data set, 80 PEER factors were generated using default settings and no covariate adjustment as per Stegle et al. using the PEER R package(4).

### eQTL Analysis

Local eQTL analysis was performed in ECLIPSE 1 and ECLIPSE 2 using the MatrixeQTL package(21). eQTL analysis for a subset of the ECLIPSE 1 subjects has been previously reported(22). All SNPs with minor allele frequency ≥5% within 1 megabase of the start and end of the gene were tested. Genotypes were imputed against the 1000 Genomes version 3 reference panel using Mach(23) and minimac(24), as previously described. FDR values were generated using the Benjamini-Hochberg method(25). A baseline eQTL analysis was performed for each dataset with adjustment only for the first three principal components of genetic ancestry calculated from the SNP data(26). We then conducted eighty analyses sequentially increasing the number of PEER factors, resulting in a total of 81 eQTL analyses for each analyzed chromosome in each data set. An additional set of analyses were performed adjusting for white blood cell differential and PEER factors.

*Retention* was defined as the percentage of significant eQTL genes from the baseline analysis that were also identified in a given PEER factor adjusted analysis for the same data set. *Replication* was defined at the level of the eQTL gene and SNP-gene pair. An eQTL gene was replicated if any association for the gene was significant at 5% FDR in the ECLIPSE 1 analysis of interest and in either the ECLIPSE 2 baseline or maximum yield PEER factor adjusted analysis. A SNP-gene pair was replicated if the same SNP-gene pair was significant at 5% FDR in ECLIPSE 1 and either the ECLIPSE 2 baseline or maximum yield PEER factor adjusted analysis.

The permutation-based significance assessment was performed by permuting the gene expression values and rerunning each eQTL analysis. For probesets in which the non-permuted minimum p-value was exceeded 15 times in the first 1,000 permutations, no additional permutation was performed. For all other probesets, a total of 10,000 permutations were performed. To correct for multiple testing, q-values were calculated from the empiric p-values of each probeset, and probesets with a q-value ≤0.05 were declared significant.

### Replication in GTEx Whole Blood Cis eQTL Results

GTEx v4 whole blood cis eQTL results were downloaded from the GTEx portal(10). All SNP-gene pairs with q-value ≤0.05 in either the ECLIPSE 1 genomewide cis eQTL baseline or max yield analysis were extracted from the GTEx results, and GTEx q-values were calculated. q-value ≤0.05 in GTEx defined replication.

### Identification of PICS SNPs and Overlap with Roadmap Epigenomic Cell-Type Enhancer Regions

The most significant SNP for each of the top 100 eQTL genes in ECLIPSE 1 was selected. We used the probabilistic identification of causal SNPs (PICS, http://www.broadinstitute.org/pubs/finemapping) approach to identify all SNPs with a ≥5% likelihood of being the causal variant for a given eQTL(11). These SNP were then queried against enhancer regions for cell types from the Roadmap Epigenomics and ENCODE projects, in the Haploreg database(12). For each gene, we summed the number of SNPs within enhancer regions multiplied by each SNP’s PICS probability. Thus, for a given gene, the SNP-enhancer overlap score could range between 0 and 1. The average overlap score across the 100 eQTL genes was used to rank cell-types. The null distribution was estimated by randomly selecting, for each of the 100 lead eQTL SNPs, three non-significant SNPs (as defined by eQTL q-value ≥ in all ECLIPSE 1 analyses) that were matched by allele frequency and distance from the transcription start site. Cell-type enhancer enrichment was then calculated for this null set and used to normalize the enrichments for the lead eQTL SNPs. The observed enrichments for immune-related cells were compared to the non-immune cells using the Mann-Whitney test.

## Acknowledgements

We acknowledge Dr. Nan Laird for her contributions regarding statistical issues related to FDR estimation.

## Author Contributions

Study design: PJC, CPH, MO, DDS, and PP. Data acquisition: EKS, RTS. Data Analysis: PJC, MO, MP, AL, RC, EHS. Critical revision of manuscript: all authors.

